# Inhibiting *Mycobacterium tuberculosis* CoaBC by targeting a new allosteric site

**DOI:** 10.1101/870154

**Authors:** Vitor Mendes, Simon R. Green, Joanna C. Evans, Jeannine Hess, Michal Blaszczyk, Christina Spry, Owain Bryant, James Cory-Wright, Daniel S-H. Chan, Pedro H.M. Torres, Zhe Wang, Sandra O’Neill, Sebastian Damerow, John Post, Tracy Bayliss, Sasha L. Lynch, Anthony G. Coyne, Peter C. Ray, Chris Abell, Kyu Y. Rhee, Helena I. M. Boshoff, Clifton E. Barry, Valerie Mizrahi, Paul G. Wyatt, Tom L. Blundell

**Affiliations:** Department of Biochemistry, University of Cambridge, 80 Tennis Court Road, Cambridge, CB2 1GA, UK; Drug Discovery Unit, College of Life Sciences, University of Dundee, Dow Street, Dundee, DD1 5EH, Scotland, UK; MRC/NHLS/UCT Molecular Mycobacteriology Research Unit & DST/NRF Centre of Excellence for Biomedical TB Research & Wellcome Centre for Infectious Diseases Research in Africa, Institute of Infectious Disease and Molecular Medicine and Department of Pathology, Faculty of Health Sciences, University of Cape Town, Anzio Road, Observatory 7925, South Africa; Department of Chemistry, University of Cambridge, Lensfield Road, Cambridge, CB2 1EW, UK; Division of Infectious Diseases, Weill Department of Medicine, Weill Cornell Medical College, New York, NY 10065, USA; Tuberculosis Research Section, Laboratory of Clinical Immunology and Microbiology, National Institute of Allergy and Infectious Disease, National Institutes of Health, 9000 Rockville Pike, Bethesda, Maryland 20892, USA

## Abstract

Coenzyme A (CoA) is a fundamental co-factor for all life, involved in numerous metabolic pathways and cellular processes, and its biosynthetic pathway has raised substantial interest as a drug target against multiple pathogens including *Mycobacterium tuberculosis*. The biosynthesis of CoA is performed in five steps, with the second and third steps being catalysed in the vast majority of prokaryotes, including *M. tuberculosis*, by a single bifunctional protein, CoaBC. Depletion of CoaBC was found to be bactericidal in *M. tuberculosis*. Here we report the first structure of a full-length CoaBC, from the model organism *Mycobacterium smegmatis*, describe how it is organised as a dodecamer and regulated by CoA thioesters. A high-throughput biochemical screen focusing on CoaB identified two inhibitors with different chemical scaffolds. Hit expansion led to the discovery of potent inhibitors of *M. tuberculosis* CoaB, which we show to bind to a novel cryptic allosteric site within CoaB.

## Introduction

Tuberculosis (TB) is the most prevalent and deadly infectious disease worldwide and remains a global epidemic. Despite the availability of treatment, this disease, caused by *Mycobacterium tuberculosis*, still claims 1.5 million lives each year ^1^. Current treatment regimens are long, which presents an obstacle for patient adherence and imposes a heavy social and economic burden on countries with a high incidence of TB. It is therefore critical to explore novel targets and find new and more effective drugs to combat this disease.

Coenzyme A (CoA) is an essential and ubiquitous cofactor involved in numerous metabolic pathways with a large number of different enzymes requiring it for their activity^2^. CoA is essential for the synthesis of phospholipids, fatty acids, polyketides, and non-ribosomal peptides, for the operation of the tricarboxylic acid cycle and in the degradation of lipids ^3^. The importance of CoA for essential post-translational modifications of proteins is also well established in both eukaryotes and prokaryotes, with various proteins post-translationally modified by thioester derivatives of CoA (acylation) or CoA itself (phosphopantetheinylation and CoAlation), while several other post-translational modifications depend indirectly on CoA through the mevalonate pathway ^4–7^. Furthermore, dephospho-CoA, an intermediate of the CoA pathway, is incorporated into some RNA transcripts during transcription initiation thereby serving as a non-canonical transcription initiating nucleotide ^8^. These RNA modifications have functional consequences and occur in both eukaryotes and bacteria ^8^. In *M. tuberculosis*, CoA plays a pivotal role in the biosynthesis of complex lipids that are crucial components of the cell wall and required for pathogenicity ^9^. It is also needed for the degradation of lipids, including cholesterol, which are the primary source of energy for this organism during infection ^10,11^. Given its ubiquitous nature, wide metabolic and functional impact of its inhibition, and lack of sequence conservation between prokaryotes and humans, the CoA pathway is therefore an attractive pathway for drug discovery for many different infectious diseases, including TB.

The biosynthesis of CoA from pantothenic acid (vitamin B_5_) is performed in five steps, sequentially catalysed by the enzymes pantothenate kinase (CoaA, also known as PanK), phosphopantothenoylcysteine synthetase (CoaB), phosphopantothenoylcysteine decarboxylase (CoaC), phosphopantetheine adenylyltransferase (CoaD) and dephospho-CoA kinase (CoaE). However, in the vast majority of prokaryotes, including *M. tuberculosis*, CoaB and CoaC are encoded by a single gene to produce a fused bifunctional enzyme (CoaBC). Transcriptional silencing of individual genes of the CoA biosynthetic pathway of this pathogen identified CoaBC as uniquely bactericidal within the CoA pathway, highlighting it as a good candidate for drug discovery ^12^.

CoaBC converts 4’-phosphopantothenate to 4’-phosphopantetheine in three steps. First, 4’-phosphopantothenate (*P*PA) reacts with CTP to form 4’-phosphopantothenoyl-CMP with the release of pyrophosphate. This intermediate subsequently reacts with cysteine to form 4’-phosphopantothenoylcysteine (*P*PC) with the release of CMP, with these two steps being catalysed by CoaB. The product of CoaB is then decarboxylated by CoaC, an enzyme of the homo-oligomeric flavin-containing decarboxylase (HFCD) protein family, to 4’-phosphopantetheine. X-ray crystal structures have been reported for the individual CoaB and CoaC enzymes in several organisms, including a structure of CoaB from *Mycobacterium smegmatis*, a close relative of *M .tuberculosis*. However, a structure of a full-length bifunctional CoaBC had not been determined.

Here we report the structure of the bifunctional CoaBC of *M. smegmatis* at 2.5 Å. We identify a previously unknown allosteric site in CoaB and crucially, we report the discovery of the first *M. tuberculosis* CoaBC allosteric inhibitors. Using X-ray crystallography and enzyme kinetic experiments, we define the mode of binding of one of the inhibitors and show its impact on the protein structure and function. These results further illustrate the potential of CoaBC as a novel drug target in *M. tuberculosis*.

## Results

### Overall structure of CoaBC

As the HFCD protein family of flavin-binding proteins are known to form homo-oligomers ^13^, we performed native electrospray-ionization mass spectrometry (ESI-MS) to investigate the stoichiometry of CoaBC, previously proposed to form a dodecamer ^13^. Both *M. tuberculosis* CoaBC (MtbCoaBC) (Figure S1A) and *M. smegmatis* CoaBC (MsmCoaBC) (Figure S1B) exclusively exhibited a dodecameric assembly, with no other oligomeric species observed in the spectra, which is consistent with a strong interaction between the subunits of the complex. The dodecamer of MtbCoaBC was centred around the 56+ charge state, with an observed mass of 537 kDa, while the dodecamer of MsmCoaBC was centred around the 52+ charge state, with an observed mass of 523 kDa. These masses are 1-2% higher than the expected masses of 525 and 518 kDa for MtbCoaBC and MsmCoaBC respectively, which can be attributed to the non-specific binding of solvent molecules or ions to the protein complexes under the soft ionization conditions employed.

Structures of a few proteins of the HFCD family have been determined ^14–18^. All of these structures show either a homo-trimeric or homo-dodecameric arrangement of the flavin-containing Rossmann-fold with trimers forming at each of the vertices of the tetrahedron in the case of a dodecameric arrangement ^15^. However, all of these proteins, unlike CoaBC, contain only a single functional domain. We solved the structure of MsmCoaBC (PDB: 6TGV) at 2.5 Å resolution (Figure 1A), in the presence of CTP and FMN (Figure 1B, S2A and S2B), using crystals belonging to the H3_2_ space group with an asymmetric unit containing four protomers forming two CoaBC dimers. Data collection and refinement statistics are summarised in (Table S1). The final model (residues 2-412) covers both CoaC and CoaB, but densities for several residues in three loop regions in CoaB are not observed (residues 290-298, 336-342, 363-376). Nevertheless, all these residues except for 375 and 376, can be seen in the MsmCoaB X-ray crystal structure (PDB: 6TH2) that we also solved in this work at 1.8 Å. The N-terminal CoaC of MsmCoaBC (residues 1-179) forms the same type of dodecameric arrangement seen in other HFCD family proteins, such as the peptidyl-cysteine decarboxylase EpiD ^15^, and it sits at the core of the dodecamer (Figure 1A and 1C) with the two domains connected through a small loop region (residues 180-189) that tightly interacts with both. The active site of CoaC sits at the interface between two protomers of one CoaC trimer and a protomer of an adjacent CoaC trimer with the FMN site facing inwards towards the hollow centre of the dodecamer (Figure 2A). A previously described flexible flap that encloses the reaction intermediate bound to *Arabidopsis thaliana* CoaC ^19^ is also observed in some of the protomers, but in an open conformation (Figure 1B).

**Figure 1:**
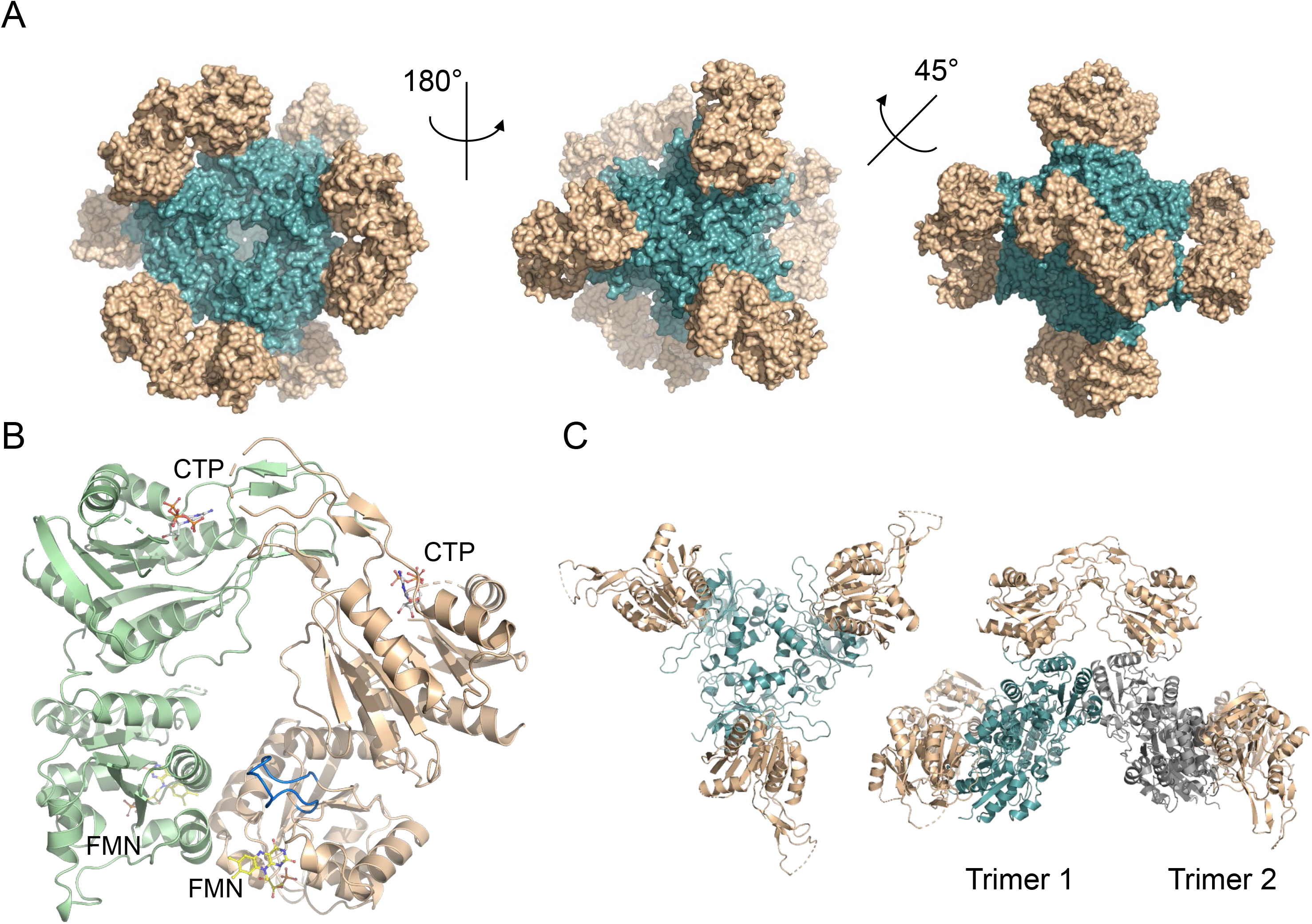
X-ray crystal structure of FMN and CTP bound MsmCoaBC. (A) Full aspect of the dodecameric CoaBC with CoaC represented in teal and CoaB in gold. (B) View of a CoaBC dimer with FMN and CTP shown. Each protomer is coloured differently. The CoaC active site flexible flap is highlighted in blue. (C) In the left panel, a CoaBC trimer is shown with the CoaC coloured in teal and CoaB in gold. On the right panel dimerization of two CoaBC trimers is shown with CoaC coloured in teal or grey for different trimers. Each CoaB forms a dimer with protomers from different trimers.

**Figure 2:**
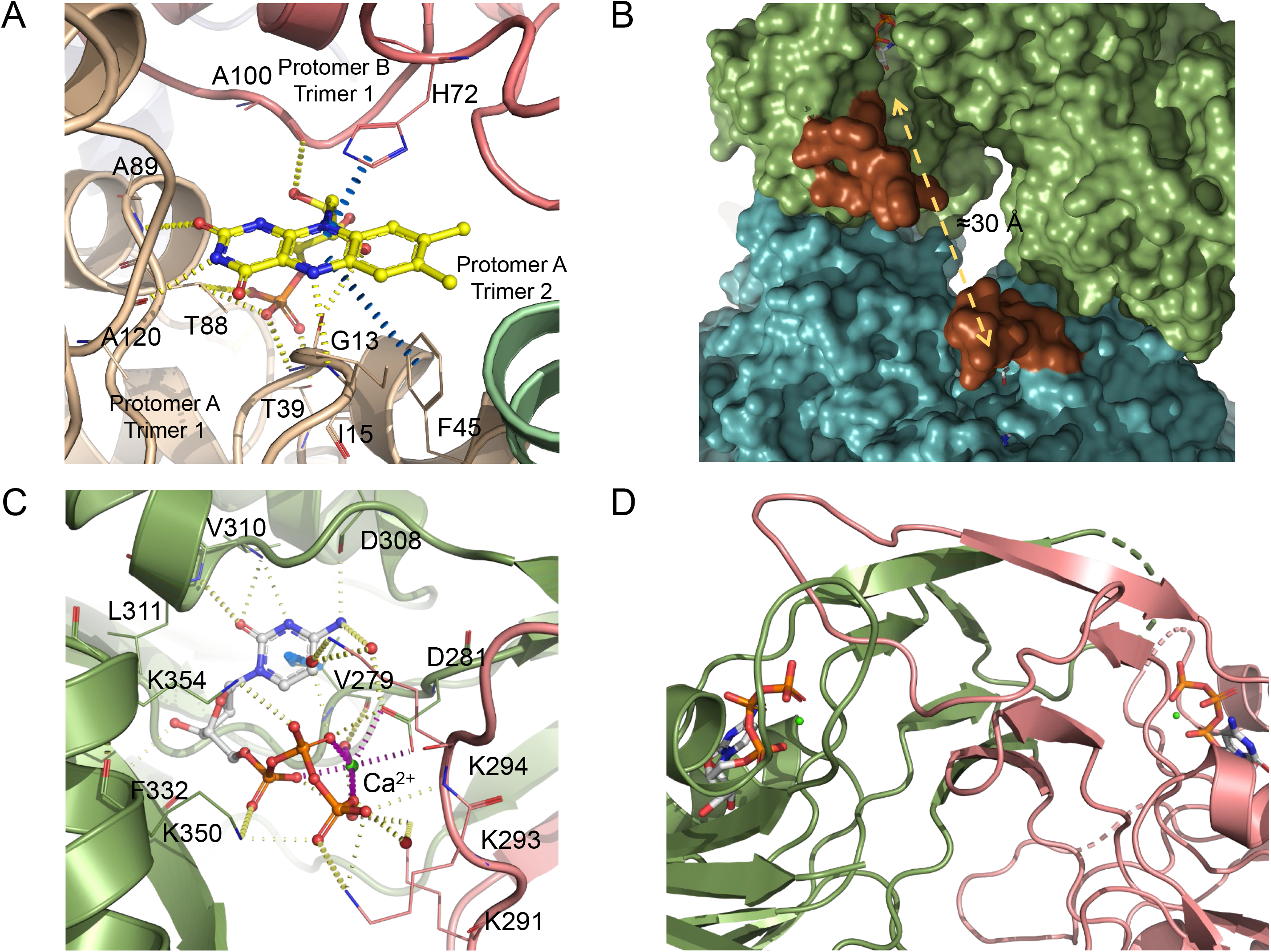
Detailed view of MsmCoaBC active sites and MsmCoaB dimerisation interface. (A) View of CoaC active site with FMN bound. The active site sits between two protomers of one trimer (gold and pink) and a third protomer from an adjacent trimer (green). Hydrogen bonds are depicted in yellow and π-interactions are in blue. (B) Superposition of a CoaB crystal structure in green, with full length CoaBC (teal) showing the active site flaps (brown) of the CoaB and CoaC enzymes. (C) Detailed view of the CTP binding site. Cartoon and residues belonging to each protomer are coloured differently. Hydrogen bonds and π-interactions are coloured as in B. Important waters are represented as red spheres and calcium as a green sphere. Calcium coordination is depicted in purple. (D) CoaB dimerization interface. Each protomer is coloured as in C. (D)

The C-terminal CoaB of MsmCoaBC also displays a Rossmann fold consistent with several other CoaB structures solved previously, including both the eukaryotic form, in which CoaB exists as an individual polypeptide, and the bacterial form where CoaB is typically fused with CoaC ^20–22^. Each CoaB of MsmCoaBC (residues 190-414) dimerises with a CoaB belonging to an adjacent trimer (Figure 1C). The full protein resembles a tetrahedron with CoaB dimers positioned at the six edges and CoaC trimers at the four vertices (Figure 1A).

The shortest distance between a pair of CoaB and CoaC active sites is ~30 Å (Figure 2B). Nevertheless, a flexible loop (residues 362-377) that covers the 4’-phosphopantothenate site, when this substrate binds to the enzyme ^20^, can be seen in our MsmCoaB structure, extending away from the active site. A superposition of our MsmCoaB dimer structure with MsmCoaBC shows the loop extending towards the CoaC active site (Figure 2B). This long loop (15-16 amino acids) is present in all CoaBCs (Figure S3) and it is possible that it helps channelling the substrate from the CoaB to the CoaC active site.

The small differences (RMSD = 1.147) in overall structure of CoaB dimers in the full length MsmCoaBC and the MsmCoaB crystal structure solved at 1.8 Å (Figure S4) can be attributed to artefacts of crystal packing. Similarly, the CoaC structure does not seem to differ between full length MsmCoaBC and the available individual CoaC structures. However, when MsmCoaB (residues 186-414) is expressed alone, the protein does not dimerise in solution and is inactive (not shown). This contrasts with *E. coli* CoaB, which still dimerises and is functional when expressed on its own without the N-terminal CoaC ^23,24^. The CoaB dimer interface is mostly conserved, but there are clear differences in the dimerisation region between MsmCoaB and *E. coli* CoaB that could help to explain the different observed oligomerisation patterns (Figure S3). The absence of dimerisation for the MsmCoaB when expressed alone suggests that the interactions between CoaC and CoaB in *M. smegmatis,* and likely all other *Mycobacteriaceae*, are fundamental for CoaB dimerisation and activity. This idea is reinforced by the fact that the residues located at the interface of the two enzymes (CoaB and CoaC) are well conserved in all *Mycobacteriaceae* and somewhat conserved in the sub-order *Corynebacterineae,* but not outside of this group (Figure S3).

The CoaB dimerisation region forms a β-sandwich composed of eight anti-parallel β-strands, related by 2-fold symmetry, that contacts with the active site (Figure 2C and 2D). Comparison of the MsmCoaB with human CoaB reveals that the human and many other eukaryotic CoaBs ^21^ possess two extra α-helices and β-strands involved in the dimerisation interface that help stabilise the dimer in the absence of CoaC (Figure S5). The CoaB active site is enclosed by a loop that extends from the opposing protomer and is observed for the first time in this work. This loop contains a motif “K-X-K-K”, which is widely conserved in bacteria (Figure S3), with few exceptions, and each lysine either interacts directly with the triphosphate group of CTP or through highly coordinated waters (Figure 2C). Also observed for the first time is the coordination of a cation by the triphosphate group and D281 (Figure 2C and S6). While magnesium or manganese are the favoured cations for CoaB activity ^25^, calcium is observed in our structures instead, due to the high concentration present in the crystallization condition.

### CoaBC is regulated by CoA thioesters

It is known that CoA biosynthesis is regulated, in many organisms, at the first step of the pathway, which is catalysed by the enzyme CoaA ^3^. *M. tuberculosis* and many other mycobacteria possess a CoaA (type I PanK) as well as CoaX (type III PanK). However, only the type I PanK seems to be active based on studies in *M. tuberculosis* ^26^. CoA and its thioesters competitively inhibit *E. coli* CoaA by binding to the ATP site, with CoA being the strongest regulator ^27,28^. Nevertheless, at physiologically relevant levels of CoA there is only a low level inhibition of CoaA ^28^. It is also known that *M. tuberculosis* CoaD, the enzyme that catalyses the fourth step of the pathway, is competitively inhibited by CoA and its product dephospho-CoA ^29,30^. However, nothing was known about the regulation of CoaBC in any organism. We therefore examined the effect of CoA and several of its thioesters (acetyl-CoA, malonyl-CoA and succinyl-CoA) on MtbCoaBC activity, using a coupled enzymatic assay that quantifies the release of pyrophosphate (EnzChek pyrophosphate assay). Controls were performed to assess the activity of these compounds against the two coupling enzymes and the compounds showed an absence of inhibition at the tested range of concentrations.

Inhibition of CoaB activity by CoA and acyl-CoAs was observed, with IC_50_ values ranging from 38 to 148 μM, far below the predicted intracellular concentrations of acyl-CoAs ^31^, with succinyl-CoA displaying the highest inhibition (Figure 3A and Table 1). Competition assays with the three substrates and acetyl-CoA show a competitive mode of inhibition relative to CTP and *P*PA with a *K_i_* of 22.5 and 22.4 μM respectively, and non-competitive inhibition for L-cysteine with a *K_i_* of 62.5 μM (Figure 3B and Table 2). In the absence of a crystal structure to confirm the mode of binding, these results suggest that acyl-CoAs most likely bind to the active site itself, competing directly with CTP and *P*PA. Interestingly, both acyl-CoAs, involved in fatty acid synthesis, as well as thoseinvolved in the TCA cycle, show inhibition of CoaB, with larger fatty acyl chains showing higher inhibition of CoaB (Figure 3A).

**Table 1:**
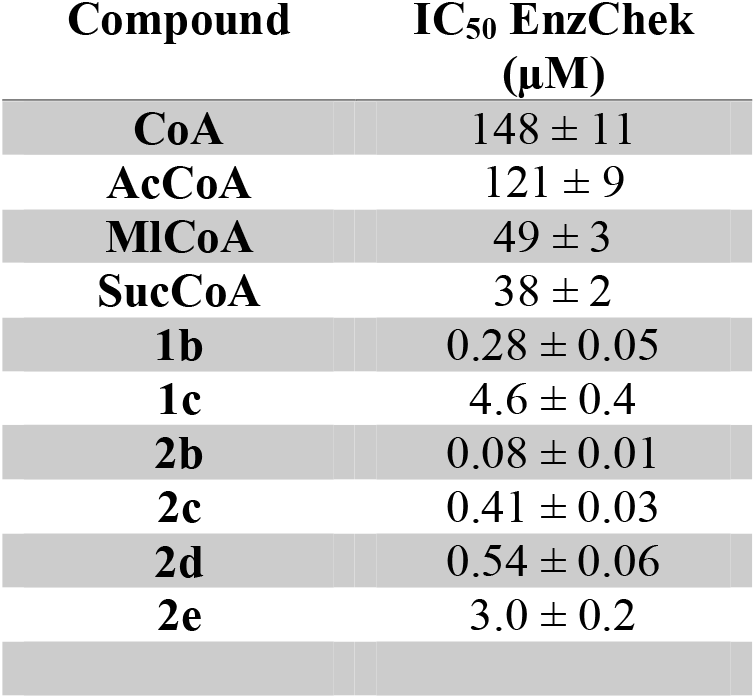
Inhibition of CoaB domain by CoA, CoA thioesters and the most potent inhibitors from of series one and two. IC_50_ values determined using the EnzChek pyrophosphate assay are shown. Error represents standard deviation with n = 3.

**Table 2:**
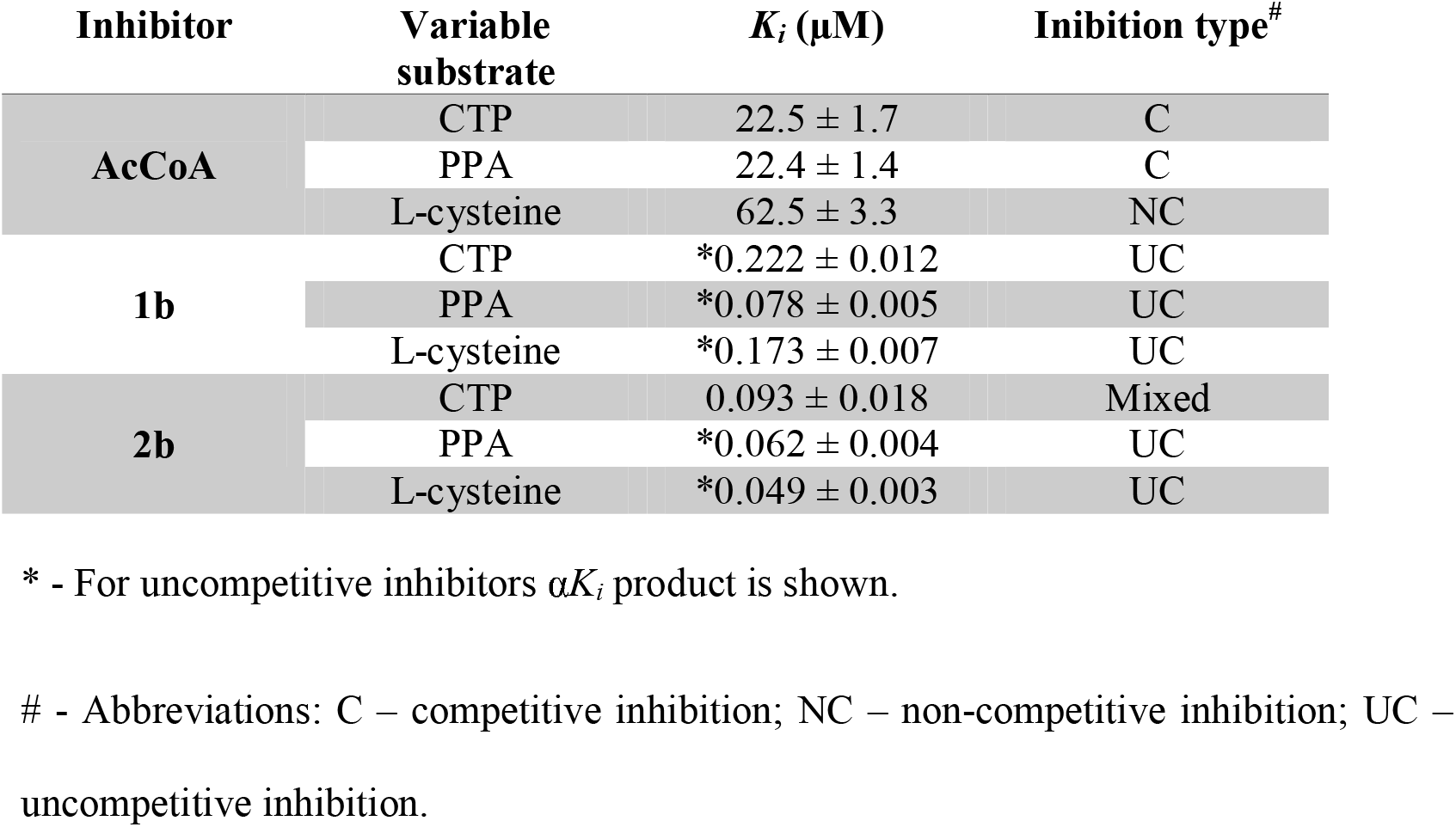
Inhibitor constants of acetyl-CoA, compound **1b** and **2b** for the three CoaB substrates. Error represents standard deviation with n = 3.

**Figure 3:**
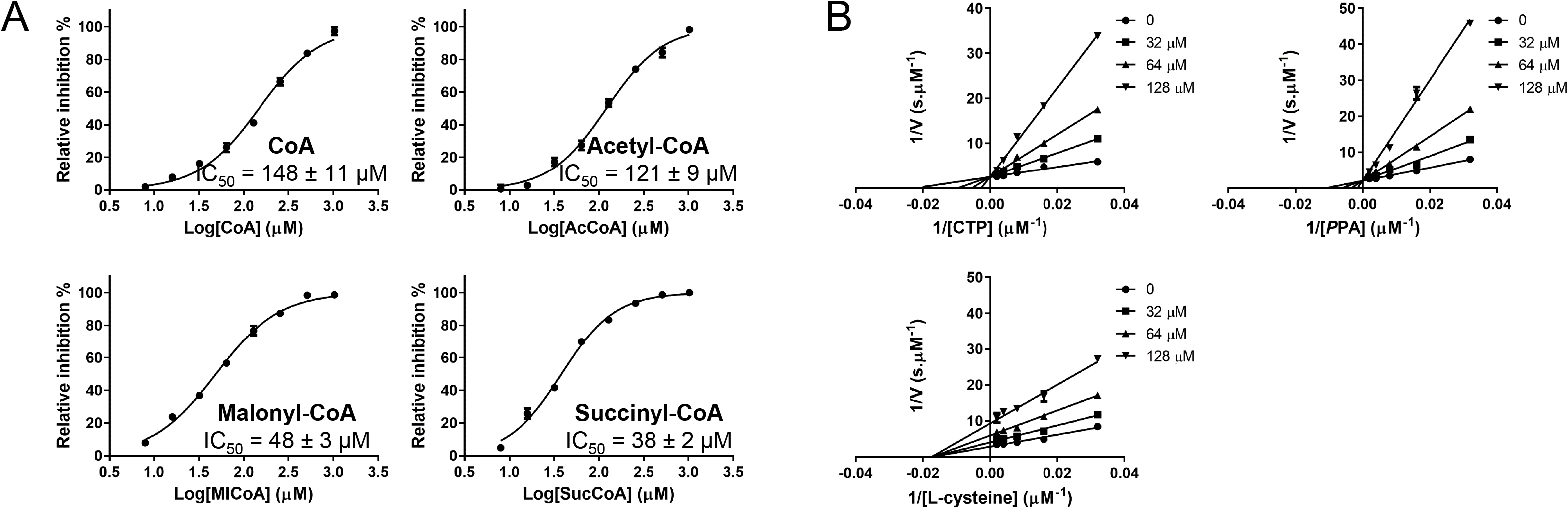
Regulation of MtbCoaBC by CoA and CoA thioesthers. (A) Inhibition of MtbCoaBC by CoA, acetyl-CoA, malonyl-CoA and succinyl-CoA. (B) Lineweaver-Burk plots showing the effect of varying the concentration of each substrate in the presence of different concentrations of acetyl-CoA. Error bars represent standard deviation with n = 3.

### Identification of CoaB inhibitors using high-throughput screening

Although the CoA biosynthetic pathway is considered an attractive target for drug discovery, CoA pathway inhibitors displaying potent whole cell activity are rare and the few CoaBC inhibitors that have been reported to date are in majority substrate mimicking ^24,32^.

In order to identify novel MtbCoaBC inhibitors, we have conducted a high-throughput screen of 215,000 small molecules targeting CoaB activity. To do this, an end-point pyrophosphate quantification assay was used (Biomol Green). The most potent hits identified were compounds **1a** and **2a** with IC_50_ values of 9 and 3.1 μM respectively (Table S2), originating from two different but related chemical scaffolds. A search was then performed for commercially available analogues. Testing of analogues of the initial hits resulted in the identification of more potent compounds with sub-micromolar IC_50_ values (Table 1, Figure 4A, Table S2 and Figure S7). Of these, compounds **1b** and **2b** (Figure 4A and Table 1), with IC_50_ values of 0.28 and 0.08 μM respectively, were identified as the most potent of the two chemical series and therefore were selected for further work.

**Figure 4:**
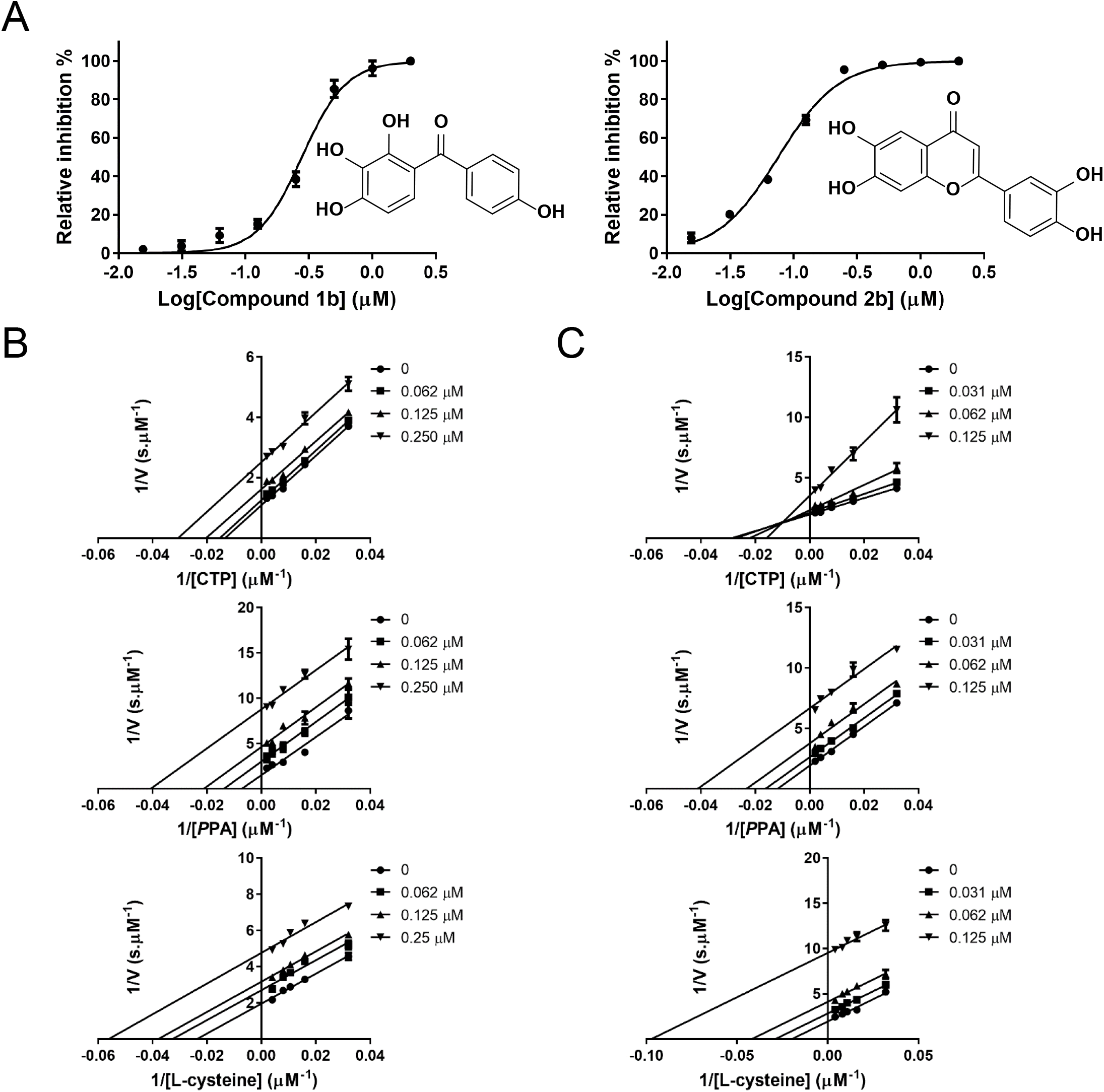
Inhibition of MtbCoaBC by compounds 1b and 2b. (A) Dose response profiles and chemical structure of compounds **1b** and **2b** is shown. (B and C) Lineweaver-Burk plots respectively showing the effect of varying concentrations of compound **1b** and **2b** in the presence of varying concentrations of CTP, *P*PA and L-cysteine. Error bars represent standard deviation with n = 3.

### Elucidation of the mode of inhibition

Following the identification of potent MtbCoaB inhibitors we went on to determine their mode of inhibition using kinetic assays. For this, the EnzChek coupled enzyme assay that measures the release of pyrophosphate was used. Control experiments were first performed to assess compound activity against the two coupling enzymes and the compounds were found to be inactive at 100 μM. The IC_50_ values for the compounds against MtbCoaB were re-determined with this assay and the values obtained were in line with the primary screening assay (Table S2).

Competition experiments were then performed between the three CoaB substrates and the two most potent compounds of each chemical series (**1b** and **2b**). Compound **1b** showed uncompetitive inhibition for all substrates with a α*K_i_* of 0.222, 0.078 and 0.173 μM respectively for CTP, *P*PA and L-cysteine (Figure 4B and Table 2), consistent with the compound binding preferentially when the three substrates are bound. Compound **2b** shows mixed inhibition relative to CTP with a *K_i_* of 0.093 μM and uncompetitive inhibition for *P*PA and L-cysteine and *P*PA with a α*K_i_* respectively of 0.062 and 0.049 μM (Figure 4C and Table 2). It is known that CoaB forms the phosphopantothenoyl-CMP intermediate in the absence of L-cysteine ^33^ and, due to spatial constraints, it is likely that cysteine can only bind at the active site after the release of pyrophosphate. The data is therefore consistent with compound **1b** preferentially binding after L-cysteine enters the active site, for the last step of catalysis and the formation of 4’-phosphopantothenoylcysteine and CMP. However, compound **2b** shows a mixed inhibition for CTP, reflecting a slightly different mechanism of action. These results obtained for both compounds suggest the existence of an allosteric site in the CoaB moiety of MtbCoaBC.

### Structural basis for inhibition of CoaB by allosteric inhibitors

In order to elucidate the binding mode of compound **1b** we used a truncation of the MsmCoaB (residues 187-414) that was previously crystallized before by others in the presence of CTP (PDB code: 4QJI) at 2.65 Å resolution. The screening for new crystallization conditions allowed us to find a new CTP containing condition that gave excellent resolution (1.8 Å). Comparison of this structure with the full length MsmCoaBC (Figure S4) showed only minor differences that can be attributed to crystal packing. Hence this crystallization system could be used to validate CoaB inhibitors binding outside of the CTP site.

MsmCoaB was co-crystallized with CTP in the absence of compound **1b** and overnight soaking of the crystals with this compound was performed. A co-crystal structure of MsmCoaB with compound **1b** was obtained and showed that the compound was bound to a site at the dimer interface of CoaB, in a deep cavity that is occluded when the compound is absent (Figure 5A-B and S8). Each CoaB dimer contains two of these sites, which are formed by a sandwich of eight β-strands and a long loop that contains the conserved “K-X-K-K” motif. This site opens to the active site and the inhibitor also contacts with D281 that is involved in the coordination of the cation (Figure 5C). The opening/closing of this cryptic allosteric site is mediated by the side chain of R207 of the opposing protomer (Figure 5D) that moves 5.5 Å at the furthest point and, to a smaller extent, by the side chain of F282 that moves 2 Å. R207 has previously been shown to be critical for the second half of the reaction catalysed by CoaB, the conversion of the 4’-phosphopantothenoyl-CMP intermediate to *P*PC, with almost no conversion of the intermediate to *P*PC detected when this arginine is mutated to glutamine ^33^. Given the position of this arginine, it is likely that it is involved in the binding of cysteine. Despite the absence of a crystal structure with cysteine, kinetic data showing uncompetitive inhibition with cysteine is consistent with this.

**Figure 5:**
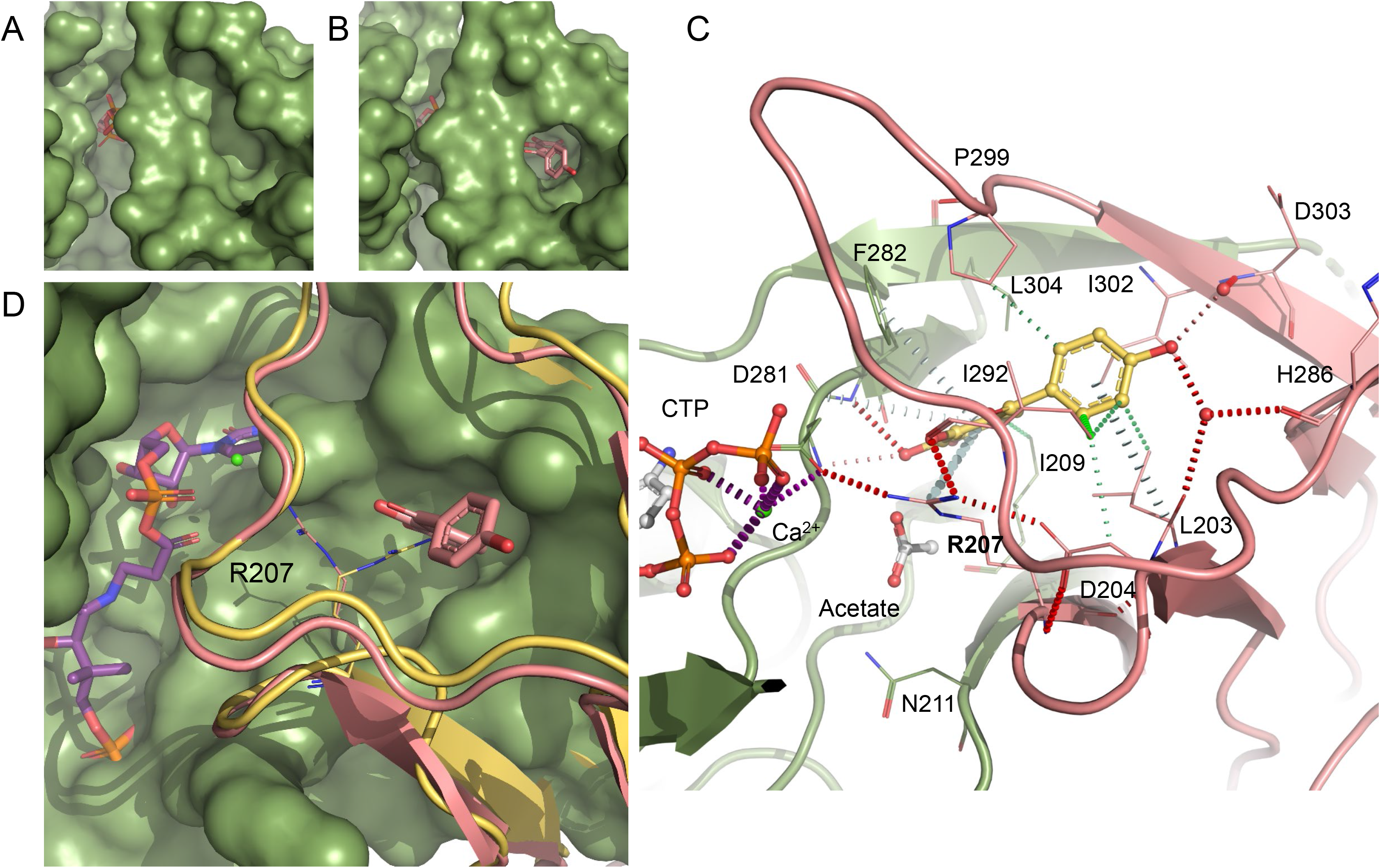
MsmCoaB X-ray structure showing the cryptic allosteric site. CoaB with the cryptic allosteric site closed (A) and opened conformation (B) with compound **1b** (pink) bound. (C) Detailed view of the allosteric site with compound **1b** (yellow) bound. The individual protomers of the CoaB dimer are coloured in green or pink. Hydrogen bonds are depicted in red, π-interactions are in grey, and hydrophobic interaction in green. Important waters are represented as red spheres and calcium as a green sphere. Calcium coordination is depicted in purple. (D) Gating mechanism of the cryptic allosteric site showing the movement of R207 with the closed conformation in yellow and the open conformation in pink. An *E. coli* structure (PDB code: 1U7Z) with the 4’-phosphopantothenoyl-CMP (purple) intermediate bound is superimposed.

This allosteric site is comprised of a large group of hydrophobic residues (I209, F282 L304 of protomer A and L203, I292, P299 and I302 of protomer B) many of which form hydrophobic interactions with compound **1b** (Figure 5C). Several π-interactions between the compound and the protein are also observed and involve D281 and F282 of protomer A and R207 of protomer B (Figure 5D). Hydrogen-bond interactions are formed with D281 and F282 of protomer A and R207 of protomer B. Water-mediated interactions are also observed for a group of residues that sit at the outer edge of the site (L203, H286 and D303) that is formed exclusively by protomer B (Figure 5C).

We propose that upon binding of L-cysteine, the R207 side chain moves towards the active site, and is likely involved in stabilizing/orienting L-cysteine to attack the phosphopantothenoyl-CMP intermediate. This movement opens the allosteric site, which allows binding of allosteric inhibitors to the newly created cavity. The allosteric inhibitors will then stabilize the enzyme in its substrate bound state with the position of R207 becoming locked by several hydrogen-bonds with the side chain of D281 of protomer A, the backbone carbonyl group of I292 and the side chain of D204 of protomer B, but also by the π-interactions with the compound (Figure 5C). The residues around this site and crucially R207 are conserved across many microorganisms, suggesting that this allosteric site is present in most, if not all bacterial CoaBCs (Figure S3 and S9A). Interestingly, even though overall sequence identity is very low between the human CoaB and MsmCoaB (22%), the human enzyme also contains an arginine equivalent to R207 and a roughly similar interface with several conserved residues, but there are stark differences in the relative position of the residues at this site between the two enzymes (Figure S9B).

While we were not able to obtain co-crystal structures with other inhibitors, *in silico* docking helped to provide a possible explanation for the structure-activity relationship observed for series one and two. The highest-scoring docking pose of compound **1b**, the most potent inhibitor of series one, was almost identical to that observed in the co-crystal structure (Figure S10A), and the analogues for which docking was performed adopted a similar binding pose. The lower activity of compound **1a** relative to compound **1b** could be explained by the loss of water-mediated hydrogen bonds (Figure 5C, S10B), while the lower activity of compound **1c** could be explained by the loss of the carbonyl group which faces a highly electropositive area of the protein (Figure S10C). Compound **2b** is predicted to form direct hydrogen bonds at the bottom of the allosteric site, similar to those formed by compound **1b**, but also to interact directly with L203 and H286, forming extra hydrogen bonds at the top of the allosteric site (Figure 6). For compound **1b** the interactions at the top of the site are water mediated (Figure 5C). This could explain the higher potency of compound **2b**. Compounds **2c** and **2d** are also predicted to form direct hydrogen bond interactions at the top of the allosteric site, but the interactions at the bottom of the site are not as favourable due to the presence of extra hydroxyl groups (Figure S10D-F). The remaining compounds in series two, which have fewer hydroxyl groups and/or hydroxyl groups in different positions, lose the ability to form hydrogen bonds, consistent with the weaker inhibitory effect observed (Table S2).

**Figure 6:**
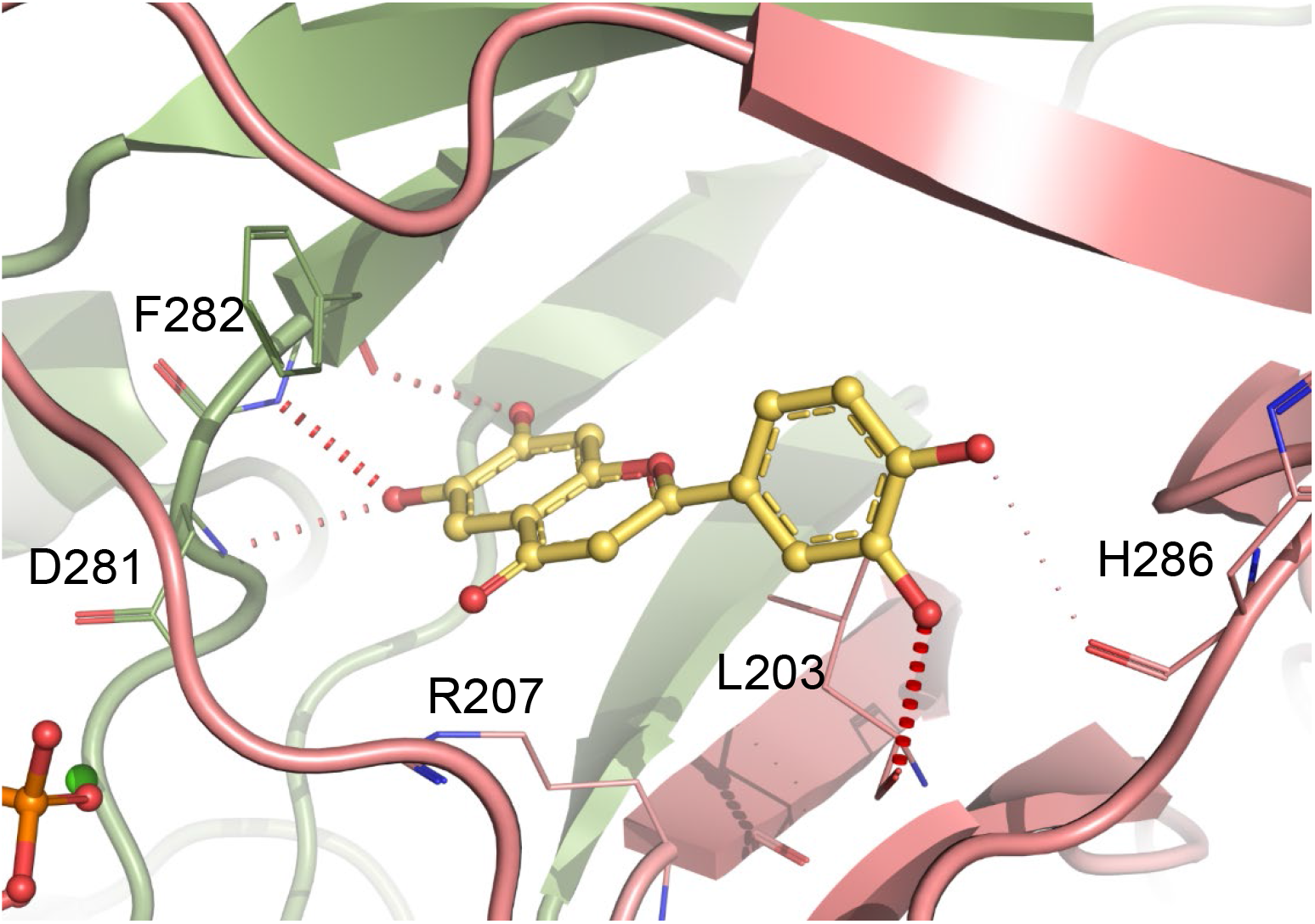
Docking of compound 2b into MsmCoaB showing the highest scoring pose. Hydrogen bonds are shown in red. The individual protomers of the CoaB dimer are either coloured in green or pink.

### Screening of CoaBC inhibitors against *M. tuberculosis*

The in vitro whole cell activity of the compounds was further evaluated by their ability to inhibit *M. tuberculosis* growth on different carbon sources. None of the compounds exhibited activity in media containing glycerol or cholesterol as the main carbon source (Table 3). We then tested whether the lack of inhibitory activity could be attributed to the presence of BSA by determining the whole cell activity of the three most potent inhibitors against *M. tuberculosis* in GAST/Fe minimal media. All the tested compounds exhibited moderate to low activity in this media with compound **2b** displaying the best activity of the three (Table 3). The observed differences in potency between the enzymatic assay and whole cell activity are likely related to low compound permeation, high efflux or metabolism.

**Table 3:** Minimum inhibitory concentration (MIC) values of CoaB inhibitors against *M. tuberculosis* H37Rv cultured in different media (μM).

| Compound | 7H9/ADC/<br>Glycerol | 7H9/<br>Cholesterol/<br>Tyloxapol | GAST/<br>Fe |
| --- | --- | --- | --- |
| **1b** | >250 | >250 | 125 |
| **1c** | >250 | ND | ND |
| **2b** | >250 | >250 | 50 |
| **2c** | >250 | >250 | 125 |
| **2d** | >250 | >250 | ND |
| **2e** | >250 | ND | ND |

\*ND – Not determined.

## Discussion

CoA is an essential co-factor ubiquitous across all domains of live. For many years, this pathway has been considered an attractive drug target to develop new antibiotics against a wide range of pathogens including *M. tuberculosis* ^34^. Furthermore, the recent identification of CoaBC as a key fragility point in the CoA pathway of this organism ^12^, combined with the extremely low sequence identity with the human CoaB (25%), makes this enzyme a highly attractive drug discovery target.

While a structure of an individual mycobacterial CoaB was available, we were aware that the many questions remaining at the start of this work about the organization and regulation of this bi-functional enzyme could have significant implications for drug discovery. We therefore set out to obtain a full-length structure of a mycobacterial CoaBC and we successfully solved the MsmCoaBC structure, which shares very high sequence identity with the *M. tuberculosis* orthologue (86% full-length, 84% CoaB enzyme) and hence is a valuable tool for studying *M. tuberculosis* CoaBC. The organization of CoaBC is similar to other HFCD family proteins ^15^ but unique in the sense that it contains more than one domain and highlights how the arrangement of the fused enzymes is essential for mycobacterial CoaB dimerisation and function. This fused arrangement might also help to channel the CoaB product to the CoaC active site more effectively. The human CoaB and other eukaryotic orthologues form stable dimers due to the extra dimerisation region (Figure S5), but is also known in yeast that the entire CoA pathway assembles into a metabolon centred on CoaC ^35,36^ (known as CAB3). This hints that close proximity between the different active sites of the CoA enzymes is desirable and that substrate channelling of products and substrates between different enzymes might be important in this pathway. It is not clear at this point if such an arrangement for the entire CoA pathway is also present in bacteria.

Regulation of the CoA biosynthesis pathway was known to occur for other enzymes of the pathway through feedback inhibition by CoA, but no information was available for CoaBC. We demonstrate that both CoA, as well as several CoA thioesters regulate CoaBC by inhibiting CoaB activity, and that these molecules act competitively for CTP and PPA and non-competitively for L-cysteine. This is consistent with these molecules binding to the CoaB active site but not to the L-cysteine sub-site. CoA and acyl-CoAs inhibit both CoaA and CoaD enzymes to varying extents ^28,30,37^. However, the inhibitory effect of CoA and its thioesters in the activity of these enzymes is lower when compared to what we observed in CoaBC and consequently the impact of the intracellular level of these molecules will be predominant in CoaBC. We therefore report a new and important mechanism of regulation of “*de novo”* CoA biosynthesis, mediated by the action of CoA thioesters on CoaBC. Since the reported intracellular levels of these molecules ^31^ are normally above the observed IC_50,_ the activity of CoaBC is highly inhibited. This correlates well with previous work showing that “de novo” CoA biosynthesis closely matches dilution due to cell division ^38^. However, the data for intracellular concentrations of CoA and CoA thioesters, as well as CoA half-life was not obtained for mycobacteria, and both interspecies differences along with variations in growth conditions may affect these conclusions.

Although the CoA pathway and CoaBC have been the subject of many drug discovery efforts, few non-substrate-mimicking inhibitors of CoaBC have been reported ^24^. Our work identifies two related chemical scaffolds that potently inhibit the activity of the CoaB moiety of MtbCoaBC through a new cryptic allosteric site that sits in the dimer interface region of the CoaB enzyme. This site is closed in the CTP-bound structure, by the side chain of R207 a residue known to be involved in the second and final step of the reaction catalysed by CoaB – the conversion of the 4’-phosphopantothenoyl-CMP intermediate to *P*PC ^33^. Considering the role of this residue in the final step of product formation and that compound **1b** shows uncompetitive inhibition relative to all CoaB substrates, we propose that the opening of this site occurs upon binding of the final substrate L-cysteine. Currently it is not clear whether this new allosteric site is exploited by a natural ligand, as we were unable to identify such a biomolecule. Nevertheless, the conservation of residues at this site, across a variety of bacteria, indicates that this feature might be common to many, if not all, bacterial CoaBs.

Drug discovery against *M. tuberculosis* is rich in examples of compounds with potent activity against an essential enzyme but with a complete lack of whole cell activity due to the impermeable cell wall of this organism, efflux pumps, target modification enzymes and extensive capacity to metabolise compounds ^39^. The modest vitro whole cell activity displayed by the CoaB inhibitors reported in this work, may relate to any of these issues. Nevertheless, the biochemical and structural data described herein further validate CoaBC as a promising novel anti-tubercular drug target by showing a new allosteric site that can be targeted by potent inhibitors.

## Materials and methods

### Cloning and protein purification

*M. tuberculosis* and *M. smegmatis coaBC* genes were amplified from genomic DNA of *M. tuberculosis* H37Rv strain, obtained from ATCC (ATCC25618D-2) and genomic DNA of *M. smegmatis* mc^2^ 155, and cloned into a pET28a vector (Novagen), modified to include an N-terminal 6xHis-SUMO tag. The *M. smegmatis coaB* construct was obtained from the Seattle Structure Genomics Center for Infectious Disease. The same protein purification protocol was used for both *M. tuberculosis* and *M. smegmatis* CoaBC constructs.

*E. coli* BL21(DE3) containing pET28aSUMO-CoaBC was grown in 2XYT media at 37°C until an O.D._600_ = 0.6. IPTG was then added to a final concentration of 0.5 mM and the temperature changed to 18 °C for 18-20h. Cells were then harvested by centrifugation, re-suspended in 50 mM TRIS pH 8.0, 250 mM NaCl, 20% (w/v) glycerol, 20 mM imidazole, 5 mM MgCl_2_ with protease inhibitors tablets (Roche) and DNAseI (Sigma). Cells were lysed with an Emulsiflex (Avestin) and the resultant cell lysate was centrifuged at 27000 *g* for 30 min to remove cell debris. Recombinant CoaBCs were purified with a HiTrap IMAC Sepharose FF column (GE-Healthcare), equilibrated with 50 mM TRIS pH 8.0, 250 mM NaCl, 20% (w/v) glycerol and 20mM Imidazole. Elution was performed in the same buffer with 500mM Imidazole. Protein was dialysed in 25 mM TRIS pH 8 and 150 mM NaCl and the SUMO tag was cleaved overnight at 4 °C by adding Ulp1 Protease at a 1:100 ratio. CoaBC was concentrated and loaded on a Superdex 200 column equilibrated with 25 mM TRIS pH 8.0, 150 mM NaCl. Fraction purity was determined by SDS-page and the purest fractions were pooled, concentrated to ~10 mg.mL^−1^ for MtbCoaBC and 30 mg.mL^−1^ for MsmCoaBC, flash frozen in liquid nitrogen and stored at −80 °C.

*E. coli* BL21(DE3) containing the *M. smegmatis* CoaB construct with a N-terminal non-cleavable 6xHis tag was grown and harvested as above and re-suspended in 20 mM HEPES pH 7.0, 500 mM NaCl, 20 mM imidazole, 5 mM MgCl_2_ with protease inhibitors tablets (Roche) and DNAseI (Sigma). Cells were lysed with an Emulsiflex (Avestin) and cell lysate was centrifuged at 27000 *g* for 30 mins to remove cell debris. Recombinant *M. smegmatis* CoaB was purified with a HiTrap IMAC Sepharose FF column (GE-Healthcare), equilibrated with 20 mM HEPES pH 7.0, 500 mM NaCl and 20 mM imidazole. Elution was carried out in the same buffer with 500 mM imidazole. Protein was concentrated and loaded on a Superdex 200 column equilibrated with 20mM HEPES pH 7.0 and 500 mM NaCl. Fraction purity was assessed by SDS-page and the purest fractions were pooled concentrated to 22 mg. mL^−1^, flash frozen in liquid nitrogen and stored at −80 °C.

### Native mass spectrometry

Spectra were recorded on a Synapt HDMS mass spectrometer (Waters) modified for studying high masses. MtCoaBC and MsCoaBC were exchanged into NH_4_OAc (500 mM, pH 7.0) solution using Micro Bio-Spin 6 chromatography columns (Bio-Rad). A sample volume of 2.5 μL was injected into a borosilicate emitter (Thermo Scientific) for sampling. Instrument conditions were optimized to enhance ion desolvation while minimizing dissociation of macromolecular complexes. Typical conditions were capillary voltage 1.8–2.0 kV, sample cone voltage 100 V, extractor cone voltage 1 V, trap collision voltage 60 V, transfer collision voltage 60 V, source temperature 20 °C, backing pressure 5 mbar, trap pressure 3–4 × 10^−2^ mbar, IMS (N_2_) pressure 5–6 × 10^−1^ mbar and TOF pressure 7–8 × 10^−7^ mbar. Spectra were calibrated externally using cesium iodide. Data acquisition and processing were performed using MassLynx 4.1 (Waters).

### Crystallization

For both full length *M. smegmatis* CoaBC and CoaB alone, the crystallization screens and optimization were performed at 18 °C using the sitting-drop vapour diffusion method. For CoaBC 300 nL of pure protein at 30 mg.mL^−1^, pre-incubated with 3 mM CTP and 10 mM MgCl_2_, was mixed in 1:1 and 1:2 (protein to reservoir) ratio with well solution using a mosquito robot (TTP labtech). Initial conditions were obtained in the Classics lite crystallization screen (Qiagen), solution 1. Crystals obtained in this condition diffracted poorly, therefore several rounds of optimization were performed. The final optimised condition consisted of 0.1 M BisTris pH 6.5, 10 mM CoCl_2_ 0.8 M 1,6-hexanediol. Crystals appeared after three days in both conditions. A cryogenic solution was prepared by adding ethylene glycol up to 30% (v/v) to the mother liquor. Crystals were briefly transferred to this solution, flash frozen in liquid nitrogen and stored for data collection.

For MsmCoaB, 200 nL of pure protein at 22-24 mg.mL^−1^ with 10 mM CTP was mixed in 1:1 ratio with well solution using a Phoenix robot (Art Robbins). Crystals were obtained in Wizards classics III&IV (Rigaku) solution G4 consisting of 20% (w/v) PEG 8000, 0.1 M MES pH 6.0 and 0.2 M calcium acetate. Crystals appeared after 2 days.

To obtain ligand-bound structures, soaking was performed condition using the hanging-drop vapour-diffusion method as follows: 2 μL of a solution containing 20% (w/v) PEG 8000, 0.1M MES pH 6.0, 0.2 M calcium acetate, 0.25 M NaCl 10% (v/v) DMSO and 1-5 mM inhibitors was left to equilibrate against 500 μL of reservoir solution for 3 days. Crystals were then transferred to the pre-equilibrated drops and incubated for 24 h. A cryogenic solution was prepared by adding 2-methyl-2,4-pentanediol up to 25% (v/v) to mother liquor. Crystals were briefly transferred to this solution, flash frozen in liquid nitrogen and stored for data collection.

### Data collection and processing

The data sets were collected at stations I02 and I03 at Diamond Light Source (Oxford, UK). The diffraction images were processed with AutoPROC ^40^ using XDS ^41^ for indexing and integration with AIMLESS ^42^ and TRUNCATE ^43^ from CCP4 Suite ^44^ for data reduction, scaling and calculation of structure factor amplitudes and intensity statistics.

### Structure solution and refinement

MsmCoaB and MsmCoaBC structures were solved by molecular replacement using PHASER ^45^ from the PHENIX software package ^46^. For MsmCoaB, the atomic coordinates of MsmCoaB structure (PDB entry 4QJI) were used as a search model. Ligand bound structures were solved using our highest resolution MsmCoaB apo form structure (PDB entry 6TH2). For MsmCoaBC, atomic coordinates of *Arabidopsis thaliana* CoaC (PDB entry 1MVL) ^19^ and our highest resolution CoaB structure (PDB entry 6TH2) were used as search models. Model building was done with Coot ^47^ and refinement was performed in PHENIX ^46^. Structure validation was performed using Coot and PHENIX tools ^46,47^. All figures were prepared using Pymol (The PyMOL Molecular Graphics System, Version 2.0 Schrödinger, LLC.) and ligand interactions calculated with Arpeggio ^48^.

### High-throughput screening

Potential inhibitors of CoaBC were assessed at room temperature using a PHERAstar microplate reader (BMG Labtech). Pyrophosphate produced by CoaB was converted to two molecules of inorganic phosphate using a pyrophosphatase. Phosphate was then detected using the BIOMOL® Green reagent (Enzo Life Sciences), which when bound to phosphate absorbs light at 650 nm. An end-point assay was carried out in clear, flat-bottom, polystyrene, 384-well plates (Greiner) in an 50 μl reaction volume containing 100 mM TRIS, pH 7.6, 1 mM MgCl_2_, 1 mM TCEP, 0.03 U/mL pyrophosphatase, 2 μM CTP, 40 μM L-cysteine, 30 μM *P*PA and 30 nM MtbCoaBC. Assays were performed by adding 25 μL of a 2-times concentrated reaction mixture containing all components with the exception of the enzymes to all wells, and the reactions started by adding 25 μL of a 2-times concentrated enzyme mixture. The reaction was carried out for 2 h at room temperature, before 50 μL of BIOMOL® Green reagent was added and incubated for a further 20 min prior to reading.

#### Inorganic pyrophosphatase-purine nucleoside phosphorylase PNP-PPIase assay

The commercially available EnzChek pyrophosphate assay kit (E-6645) (Life Technologies) was used for this assay. The final reaction composition used was 0.03 U/mL inorganic pyrophosphatase, 1 U/mL purine nucleoside phosphorylase, 1 mM MgCl_2_, 200 μM MESG, 100 mM TRIS pH 7.5, 1 mM TCEP, 32 nM MtbCoaBC, 125 μM CTP, 125 μM *P*PA, 500 μM L-cysteine, and various concentrations of compounds being tested for inhibition, all prepared from DMSO stock solutions (compounds of series one and two) or water (CoA and CoA thioesters). Assays were performed on either a CLARIOstar or PHERAstar microplate reader (BMG Labtech) in 96-well plates (Greiner). A substrate mixture containing the substrates and the inhibitor was pre-incubated at 25 °C for 10 min. An enzyme solution was prepared and separately pre-incubated at 25 °C for 10 min. The reaction was initiated by the addition of the substrates to the solution containing the enzyme to a final volume of 75 μL. Enzymatic activity was monitored by following the absorbance at 360 nm for 30 min (100 cycles/20 s each cycle). Assays were performed in triplicates, including a negative control (lacking *P*PA) and a positive control (lacking inhibitor).

Competition assays were performed using the same conditions but with variable substrate concentrations (31.25 μM, 62.5 μM, 125 μM, 250 μM and 500 μM for CTP and *P*PA, 31.25 μM, 62.5 μM, 93.75 μM 125 μM and 250 μM for L-cysteine).

### *M. tuberculosis* strains and growth conditions

MIC determination for *M. tuberculosis* H37RvMA was performed as previously described ^49^ in the following media: 7H9/ADC/glycerol (4.7 g/L Difco Middlebrook 7H9 base, 100mL/L Middlebrook albumin (BSA)-dextrose-catalase (ADC) Difco Middlebrook, 0.2% glycerol and 0.05% Tween-80), 7H9/Cholesterol/Tyloxapol (4.7 g/L 7H9 base, 0.81 g/L NaCl, 24 mg/L cholesterol, 5 g/L BSA fraction V and 0.05% Tyloxapol) and GAST/Fe (0.3 g/L of Bacto Casitone (Difco), 4.0 g/L of dibasic potassium phosphate, 2.0 g/L of citric acid, 1.0 g/L of L-alanine, 1.2 g/L of magnesium chloride hexahydrate, 0.6 g/L of potassium sulfate, 2.0 g/L of ammonium chloride, 1.80 ml/L of 10 N sodium hydroxide, 10.0 ml of glycerol 0.05% Tween 80 and 0.05 g of ferric ammonium citrate adjusted to pH 6.6).

## Acknowledgements

This work was funded by the Bill and Melinda Gates Foundation HIT-TB (OPP OPP1024021) and SHORTEN-TB (OPP1158806) (VMendes and JCE) and in part by the Intramural Research Program of NIH, NIAID (HIMB and CEB) and the South African Medical Research Council and National Research Foundation (VMizrahi). CS was funded in part by a NHMRC Overseas Biomedical Fellowship (1016357) and in part by the Bill and Melinda Gates Foundation HIT-TB (OPP OPP1024021). CoaBC screening was funded by a MRC-CinC (grant no. MC_PC_14099). TLB is funded by the Wellcome Trust (Wellcome Trust Investigator Award 200814_Z_16_Z: RG83114). The authors would like to thank the Diamond Light Source for beam-time (proposals mx9537, mx14043, mx18548), the Seattle Structural Genomics Centre for Infectious Disease for kindly providing the *M. smegmatis* CoaB plasmid and Dr Nuno Empadinhs for graciously providing the DNA of *M. smegmatis* mc^2^ 155.

## Author contributions

VMendes wrote the manuscript. VMendes designed and performed all the crystallographic experiments with the help of MB, OB and JCW. VMendes and JH designed and performed the kinetic experiments. JH synthesised 4’-phosphopantothenate. PHMT performed docking experiments. DSC performed the native mass spectrometry experiments. SG, TB, SON, SD, JP and CS developed and performed the high-throughput screening. JCE, SLL and HIMB performed the microbiology experiments on *M. tuberculosis* H37Rv. VMendes, JCE, SG, AGC, PCR, KYR, CA, HIMB, CEB, VMizrahi, PGW and TLB managed the project. All authors approved the manuscript.

## Accession numbers

Coordinates and structure factors related to this work have been deposited in the PDB with accession numbers: **6TGV**, **6TH2** and **6THC**.

## References

1. World Health, O. Global tuberculosis report 2019, (World Health Organization, Geneva, 2019).

2. Strauss, E. Coenzyme A Biosynthesis and Enzymology. Comprehensive Natural Products Ii: Chemistry and Biology, Vol 7: Cofactors, 351–410 (2010).

3. Leonardi, R. & Jackowski, S. Biosynthesis of Pantothenic Acid and Coenzyme A. EcoSal Plus 2 (2007).

4. Tsuchiya, Y. et al. Protein CoAlation and antioxidant function of coenzyme A in prokaryotic cells. Biochem J 475, 1909–1937 (2018).

5. Choudhary, C. et al. Lysine acetylation targets protein complexes and co-regulates major cellular functions. Science 325, 834–40 (2009).

6. Beld, J., Sonnenschein, E.C., Vickery, C.R., Noel, J.P. & Burkart, M.D. The phosphopantetheinyl transferases: catalysis of a post-translational modification crucial for life. Nat Prod Rep 31, 61–108 (2014).

7. Wang, M. & Casey, P.J. Protein prenylation: unique fats make their mark on biology. Nat Rev Mol Cell Biol 17, 110–22 (2016).

8. Bird, J.G. et al. The mechanism of RNA 5’ capping with NAD+, NADH and desphospho-CoA. Nature 535, 444–7 (2016).

9. Marrakchi, H., Laneelle, M.A. & Daffe, M. Mycolic acids: structures, biosynthesis, and beyond. Chem Biol 21, 67–85 (2014).

10. Guerrini, V. et al. Storage lipid studies in tuberculosis reveal that foam cell biogenesis is disease-specific. PLoS Pathog 14, e1007223 (2018).

11. Peyron, P. et al. Foamy macrophages from tuberculous patients’ granulomas constitute a nutrient-rich reservoir for M. tuberculosis persistence. PLoS Pathog 4, e1000204 (2008).

12. Evans, J.C. et al. Validation of CoaBC as a Bactericidal Target in the Coenzyme A Pathway of Mycobacterium tuberculosis. ACS Infect Dis 2, 958–968 (2016).

13. Kupke, T. et al. Molecular characterization of lantibiotic-synthesizing enzyme EpiD reveals a function for bacterial Dfp proteins in coenzyme A biosynthesis. J Biol Chem 275, 31838–46 (2000).

14. White, M.D. et al. UbiX is a flavin prenyltransferase required for bacterial ubiquinone biosynthesis. Nature 522, 502–6 (2015).

15. Blaesse, M., Kupke, T., Huber, R. & Steinbacher, S. Crystal structure of the peptidyl-cysteine decarboxylase EpiD complexed with a pentapeptide substrate. EMBO J 19, 6299–310 (2000).

16. Blaesse, M., Kupke, T., Huber, R. & Steinbacher, S. Structure of MrsD, an FAD-binding protein of the HFCD family. Acta Crystallogr D Biol Crystallogr 59, 1414–21 (2003).

17. Albert, A. et al. The X-ray structure of the FMN-binding protein AtHal3 provides the structural basis for the activity of a regulatory subunit involved in signal transduction. Structure 8, 961–9 (2000).

18. Manoj, N. & Ealick, S.E. Unusual space-group pseudosymmetry in crystals of human phosphopantothenoylcysteine decarboxylase. Acta Crystallogr D Biol Crystallogr 59, 1762–6 (2003).

19. Steinbacher, S. et al. Crystal structure of the plant PPC decarboxylase AtHAL3a complexed with an ene-thiol reaction intermediate. J Mol Biol 327, 193–202 (2003).

20. Stanitzek, S., Augustin, M.A., Huber, R., Kupke, T. & Steinbacher, S. Structural basis of CTP-dependent peptide bond formation in coenzyme A biosynthesis catalyzed by Escherichia coli PPC synthetase. Structure 12, 1977–88 (2004).

21. Manoj, N., Strauss, E., Begley, T.P. & Ealick, S.E. Structure of human phosphopantothenoylcysteine synthetase at 2.3 A resolution. Structure 11, 927–36 (2003).

22. Zheng, P. et al. Crystallographic Analysis of the Catalytic Mechanism of Phosphopantothenoylcysteine Synthetase from Saccharomyces cerevisiae. J Mol Biol (2019).

23. Kupke, T. Molecular characterization of the 4’-phosphopantothenoylcysteine synthetase domain of bacterial dfp flavoproteins. J Biol Chem 277, 36137–45 (2002).

24. Chan, D.S.-H. et al. Structural insights into Escherichia coli phosphopantothenoylcysteine synthetase by native ion mobility–mass spectrometry. Biochemical Journal 476, 3125–3139 (2019).

25. Strauss, E., Kinsland, C., Ge, Y., McLafferty, F.W. & Begley, T.P. Phosphopantothenoylcysteine synthetase from Escherichia coli. Identification and characterization of the last unidentified coenzyme A biosynthetic enzyme in bacteria. J Biol Chem 276, 13513–6 (2001).

26. Awasthy, D. et al. Essentiality and functional analysis of type I and type III pantothenate kinases of Mycobacterium tuberculosis. Microbiology 156, 2691–701 (2010).

27. Song, W.J. & Jackowski, S. Kinetics and regulation of pantothenate kinase from Escherichia coli. J Biol Chem 269, 27051–8 (1994).

28. Vallari, D.S., Jackowski, S. & Rock, C.O. Regulation of pantothenate kinase by coenzyme A and its thioesters. J Biol Chem 262, 2468–71 (1987).

29. Wubben, T.J. & Mesecar, A.D. Kinetic, thermodynamic, and structural insight into the mechanism of phosphopantetheine adenylyltransferase from Mycobacterium tuberculosis. J Mol Biol 404, 202–19 (2010).

30. Miller, J.R. et al. Phosphopantetheine adenylyltransferase from Escherichia coli: investigation of the kinetic mechanism and role in regulation of coenzyme A biosynthesis. J Bacteriol 189, 8196–205 (2007).

31. Bennett, B.D. et al. Absolute metabolite concentrations and implied enzyme active site occupancy in Escherichia coli. Nat Chem Biol 5, 593–9 (2009).

32. Moolman, W.J., de Villiers, M. & Strauss, E. Recent advances in targeting coenzyme A biosynthesis and utilization for antimicrobial drug development. Biochem Soc Trans 42, 1080–6 (2014).

33. Kupke, T. Active-site residues and amino acid specificity of the bacterial 4’-phosphopantothenoylcysteine synthetase CoaB. Eur J Biochem 271, 163–72 (2004).

34. Spry, C., Kirk, K. & Saliba, K.J. Coenzyme A biosynthesis: an antimicrobial drug target. FEMS Microbiol Rev 32, 56–106 (2008).

35. Olzhausen, J., Moritz, T., Neetz, T. & Schuller, H.J. Molecular characterization of the heteromeric coenzyme A-synthesizing protein complex (CoA-SPC) in the yeast Saccharomyces cerevisiae. FEMS Yeast Res 13, 565–73 (2013).

36. Ruiz, A. et al. Moonlighting proteins Hal3 and Vhs3 form a heteromeric PPCDC with Ykl088w in yeast CoA biosynthesis. Nat Chem Biol 5, 920–8 (2009).

37. Yun, M. et al. Structural basis for the feedback regulation of Escherichia coli pantothenate kinase by coenzyme A. J Biol Chem 275, 28093–9 (2000).

38. Hartl, J., Kiefer, P., Meyer, F. & Vorholt, J.A. Longevity of major coenzymes allows minimal de novo synthesis in microorganisms. Nat Microbiol 2, 17073 (2017).

39. Nguyen, L. & Pieters, J. Mycobacterial subversion of chemotherapeutic reagents and host defense tactics: challenges in tuberculosis drug development. Annu Rev Pharmacol Toxicol 49, 427–53 (2009).

40. Vonrhein, C. et al. Data processing and analysis with the autoPROC toolbox. Acta Crystallogr D Biol Crystallogr 67, 293–302 (2011).

41. Kabsch, W. Xds. Acta Crystallogr D Biol Crystallogr 66, 125–32 (2010).

42. Evans, P.R. & Murshudov, G.N. How good are my data and what is the resolution? Acta Crystallographica Section D-Biological Crystallography 69, 1204–1214 (2013).

43. French, S. & Wilson, K. Treatment of Negative Intensity Observations. Acta Crystallographica Section A 34, 517–525 (1978).

44. Winn, M.D. et al. Overview of the CCP4 suite and current developments. Acta Crystallogr D Biol Crystallogr 67, 235–42 (2011).

45. McCoy, A.J. et al. Phaser crystallographic software. J Appl Crystallogr 40, 658–674 (2007).

46. Adams, P.D. et al. PHENIX: a comprehensive Python-based system for macromolecular structure solution. Acta Crystallogr D Biol Crystallogr 66, 213–21 (2010).

47. Emsley, P., Lohkamp, B., Scott, W.G. & Cowtan, K. Features and development of Coot. Acta Crystallogr D Biol Crystallogr 66, 486–501 (2010).

48. Jubb, H.C. et al. Arpeggio: A Web Server for Calculating and Visualising Interatomic Interactions in Protein Structures. J Mol Biol 429, 365–371 (2017).

49. Singh, V. et al. The complex mechanism of antimycobacterial action of 5-fluorouracil. Chem Biol 22, 63–75 (2015).

