## Supplemental data for "Inhibiting *Mycobacterium tuberculosis* CoaBC by targeting a new allosteric site"

**Results**

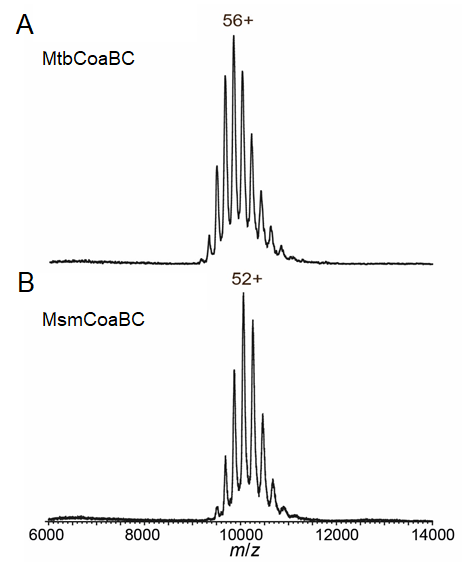

**Figure S1:** Native mass spectra of a) MtbCoaBC and b) MsmCoaBC, showing dodecameric species with charge states as indicated and masses of 537 and 523 kDa, respectively.

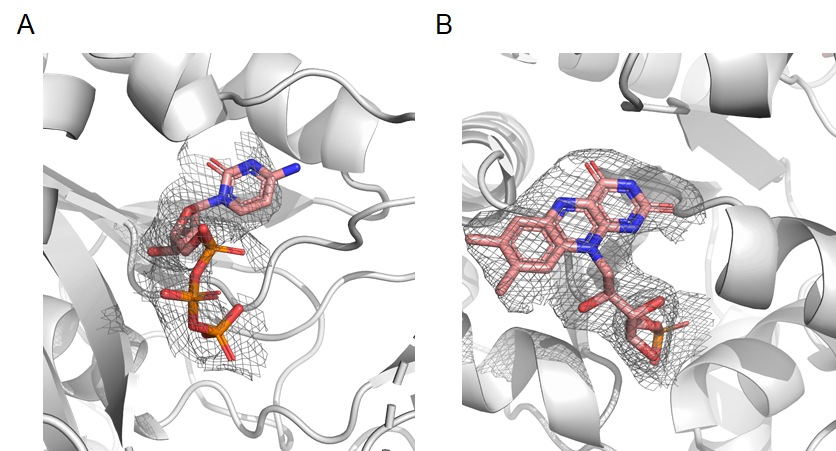

**Figure S2:** MsmCoaBC X-ray crystal structure showing Fo-Fc “Omit” maps of CTP (A) and FMN (B) contoured at 2.0 σ.

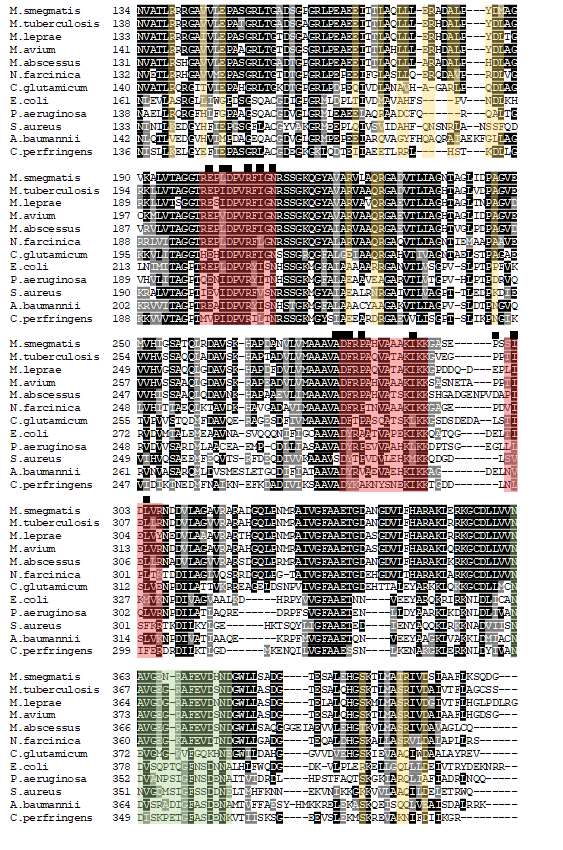

**Figure S3:** Comparison of CoaBC sequences from *Mycobacterium smegmatis*, *Mycobacterium tuberculosis*, *Mycobacterium leprae*, *Mycobacterium avium*, *Mycobacterium abscessus*, *Nocardia farcinica*, *Corynebacterium glutamicum*, *Escherichia coli*, *Pseudomonas aeruginosa*, *Staphylococcus aureus*, *Acinetobacter baumannii* and *Clostridium perfringens*. Residues that form the CoaB dimer interface are highlight in red, and those involved in the CoaB-CoaC interface in yellow. The CoaB allosteric site residues are marked with black squares above the sequences and the residues that form the CoaB loop that covers the *P*PA site are shaded in green. The residues at both the interfaces are highly conserved within *Mycobacteriaceae* pointing to shared properties of the enzyme across this group. The high conservation of allosteric site residues across diverse bacterial species is consistent with the allosteric site being a common feature of all bacterial CoaBCs. The multiple sequence alignment was performed with T-Coffee (1).

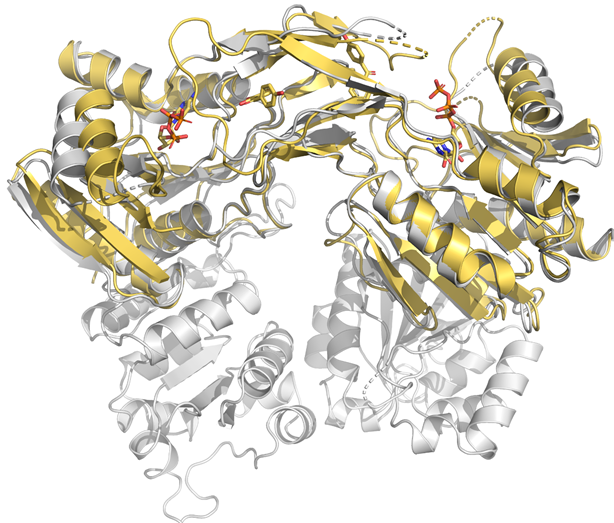

**Figure S4:** Superposition of the MsmCoaB dimer (yellow) with CTP and compound **1b** bound, with the full-length MsmCoaBC dimer (grey). The superposition shows that there are only small differences (RMSD = 1.147 Å) between the X-ray crystal structures of the individually expressed CoaB domain and the CoaB domain part of the full length CoaBC, which can be attributed to crystallographic artefacts.

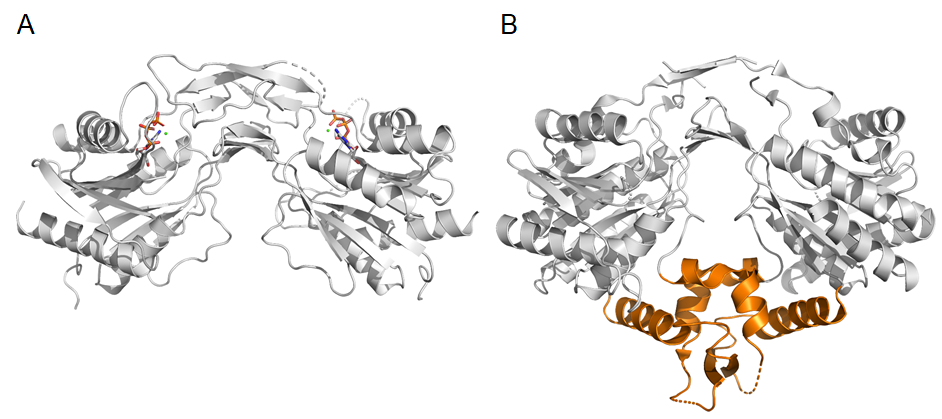

**Figure S5:** Comparison of the MsmCoaB dimer (A) and the human CoaB dimer (B). The human and other eukaryotic CoaBs have an additional two helices and β-strands (highlighted in orange) involved in dimerisation, making the dimer much more stable.

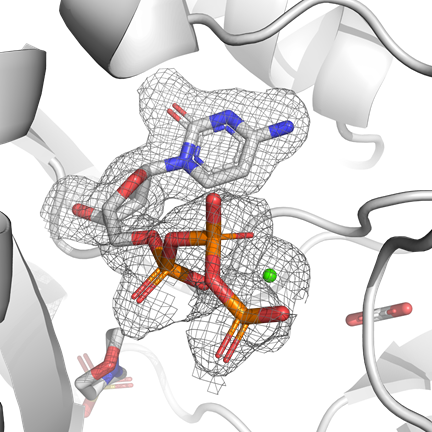

**Figure S6:** MsmCoaB X-ray crystal structure showing a Fo-Fc “Omit” map of CTP and Ca^2+^. Acetate and MES are also visible in the structure.

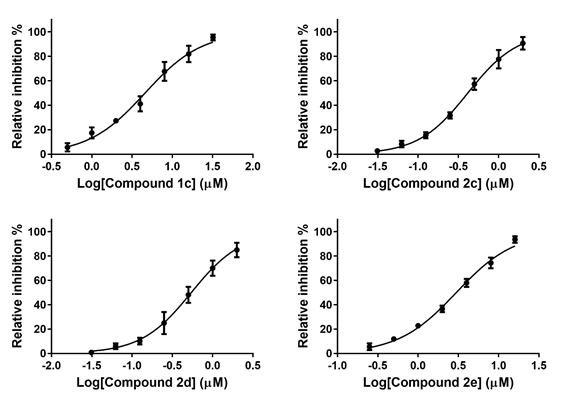

**Figure S7:** Dose response profiles for compounds **1c**, **2c**, **2d** and **2e** on CoaB activity of MtbCoaBC measured using the EnzChek pyrophosphate assay. The error bars represent standard deviation with n = 3.

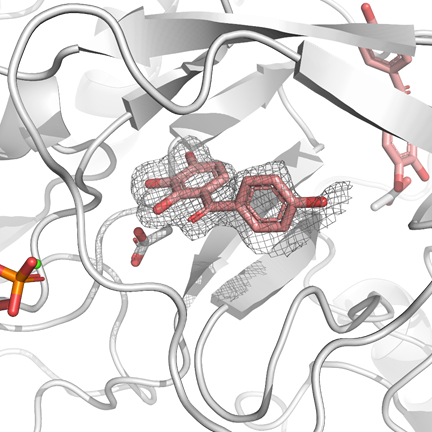

**Figure S8:** MsmCoaB X-ray crystal structure showing a Fo-Fc “Omit” map of compound **1b**. Acetate and a phosphate of CTP are also visible in the structure.

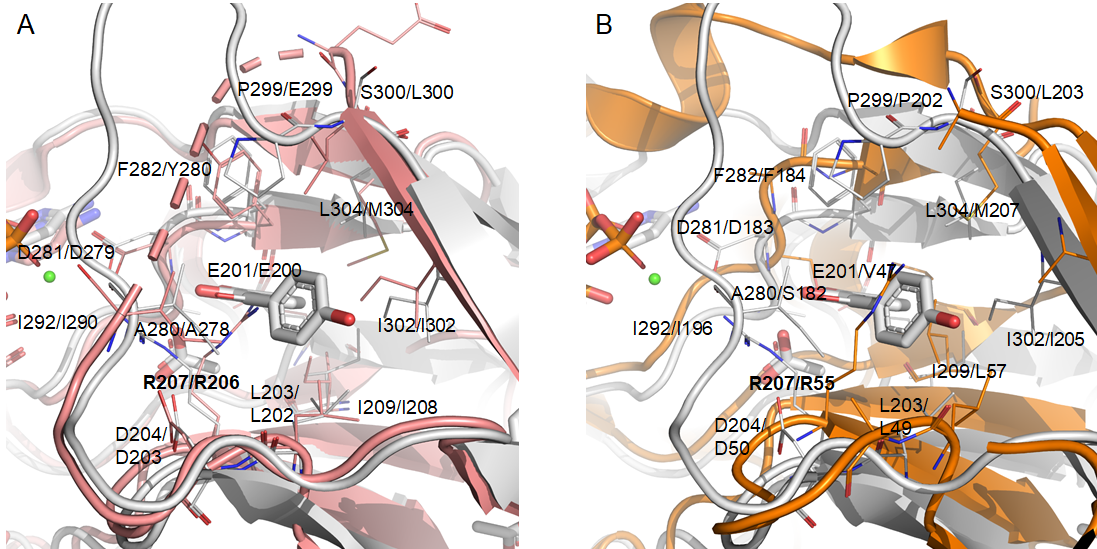

**Figure S9:** Superimposition of the MsmCoaB crystal structure in complex with compound **1b** (white) with *E. coli* CoaB (PDB: 17UZ) (pink) (A) and with human CoaB (PDB: 1P9O) (orange) (B), showing the allosteric site. Residue numbering is given for MsmCoaB first and *E. coli*/human CoaB in second. The arginine involved in the allosteric site gating and its equivalents in the *E. coli* and human CoaBs are highlighted in bold. Human CoaB residues I196 and P202 are disordered in the structure and not observed and the side chains of D50, D183, F184, L203 and I205 are also not visible. The high conservation of residues and relative positions shows that the allosteric site is also present in *E. coli* CoaBC and possibly also in the human enzyme.

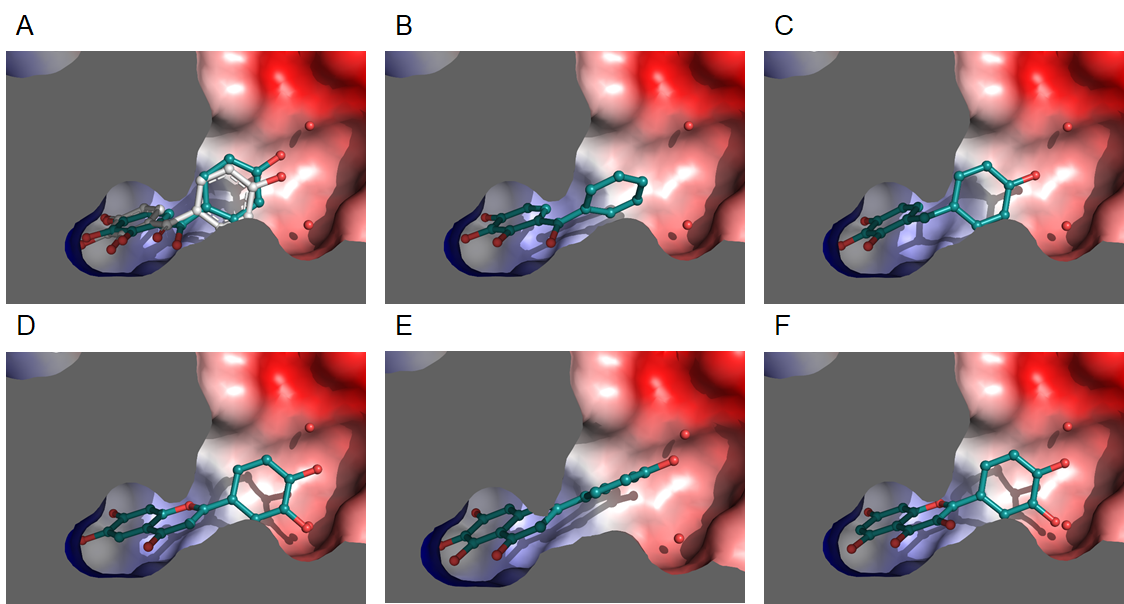

**Figure S10:** Best docking poses of compounds **1b** (A), **1a** (B), **1c** (C), **2b** (D), **2c** (E) and **2d** (F). The structure of complex of CoaB with compound **1b** was used as a receptor. A comparison of the compound **1b** crystal structure (white) and best docking pose (teal) is shown in (A). Two waters mediating interactions between compound **1b** and CoaB are shown in all figures for comparative purposes.

**Table S1:** X-ray crystallography data collection and final refinement statistics

|  | **CoaBC** | **CoaB:CTP** | **CoaB:compound 1b** |
| --- | --- | --- | --- |
| PDB ID | 6TGV | 6TH2 | 6THC |
| **Data collection*** |  |  |  |
| Space group | *H*3_2_ | *P*2_1_2_1_2_1_ | *P*2_1_2_1_2_1_ |
| Cell parameters:  a [Å]  b [Å]  c [Å]  α/β/γ [˚] | 195.11  195.11  373.92  90/90/120 | 76.50  76.84  149.05  90/90/90 | 76.01  77.37  144.35  90/90/90 |
| Resolution range [Å] | 81.80 – 2.50  (2.56 – 2.50) | 68.30 – 1.84  (1.94 – 1.84) | 77.37 – 2.03  (2.14 – 2.03) |
| No. of observations  total  unique | 1416291  (83396)  94420  (6905) | 552418  (44751)  75835  (10396) | 360758  (53437)  55496  (7959) |
| R_merge_ | 0.094 (2.987) | 0.042(0.441) | 0.093 (0.846) |
| I/σ(I) | 14.3 (1.2) | 26 (2.6) | 13.1 (2.4) |
| CC(1/2) | 0.999 (0.435) | 1 (0.878) | 0.998 (0.738) |
| Completeness [%] | 100.0 (100.0) | 99.3 (95.2) | 99.9 (99.9) |
| Multiplicity | 15.0 (12.1) | 7.3 (4.3) | 6.5 (6.7) |
| **Refinement** |  |  |  |
| Refinement program | PHENIX | PHENIX | PHENIX |
| Resolution [Å] | 81.77 – 2.50 | 50.95 – 1.84 | 72.18 – 2.31 |
| No. reflections | 94420 | 75754 | 55420 |
| R_work_/R_free_ [%] | 20.4/24.5 | 17.6/20.2 | 18.4/24.0 |
| RMS deviations |  |  |  |
| Bonds [Å] | 0.008 | 0.007 | 0.008 |
| Angles [˚] | 1.046 | 1.126 | 1.06 |
| Ramachandran |  |  |  |
| Favoured [%] | 96 | 98 | 97 |
| Outliers [%] | 0.2 | 0 | 0.1 |
| Average B-factor [Å^2^] |  |  |  |
| macromolecule | 106.0 | 37.1 | 48.8 |
| ligands | 136.7 | 43.0 | 55.1 |
| solvent | 68.0 | 40.7 | 46.4 |

* Parameters shown in brackets are for the highest resolution shell

**Table S2:** Chemical structures, and IC_50_ values for inhibition of MtbCoaB activity of all compounds in series one and two as measured by either a biomol green assay or EnzCheck assay.

| Compound | Chemical structure | IC_50_ biomol green assay (µM) | IC_50_ Enz check assay (µM) |
| --- | --- | --- | --- |
| **1a** | 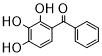 | 9 | ND |
| **1b** | 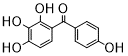 | 0.3 | 0.28 ± 0.05 |
| **1c** | 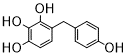 | 4.7 | 4.6 ± 0.4 |
| **1d** | 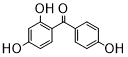 | >50 | ND |
| **1e** | 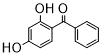 | >50 | ND |
| **1f** | 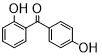 | >50 | ND |
| **1g** | 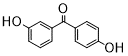 | >50 | ND |
| **2a** | 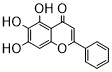 | 3.1 | ND |
| **2b** | 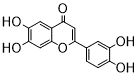 | 0.1 | 0.08 ± 0.01 |
| **2c** | 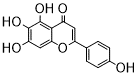 | 0.34 | 0.41 ± 0.03 |
| **2d** | 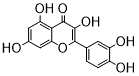 | 0.49 | 0.54 ± 0.06 |
| **2e** | 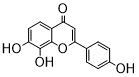 | 2.2 | 3.0 ± 0.2 |
| **2f** | 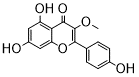 | 30 | ND |
| **2g** | 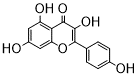 | >50 | ND |
| **2h** | 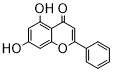 | >50 | ND |
| **2i** | 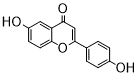 | >50 | ND |
| **2j** | 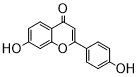 | >50 | ND |
| **2k** | 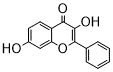 | >50 | ND |
| **2l** | 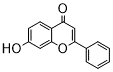 | >50 | ND |
| **2m** | 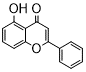 | >50 | ND |
| **2n** | 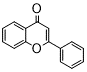 | >50 | ND |

**Materials and methods**

**Synthesis of 4’-phosphopantothenate.**

4’-Phosphopantothenate was synthesised following a literature procedure (2).

**Molecular Docking**

All evaluated compounds were generated using MarvinSketch software, ChemAxon, (http://www.chemaxon.com), saved in the PDB format and subsequently converted to .pdbqt files using AutoDockTools (3) included in the MGLTools v1.5.6 distribution. The receptor molecule used consisted of chains C and D of PDB ID: 6THC, and was also prepared using the same version of MGLTools. Polar hydrogens were added to the structure and it was saved in pdbqt format. A cubic grid box with 18 Å-long edges was manually set to loosely accommodate the allosteric binding site. Autodock VINA (4) was used to generate up to 5 poses of each ligand (num_modes=5) within a maximum energy range of 10 kcal/mol (energy_range=10) and the exhaustiveness was set to 40. The Open Drug Discovery Toolkit (ODDT) (5) was used to re-score the docked poses, using the RFScore_V3 function, trained on the PdbBind2015 dataset. To increase the robustness of the results, the above-described procedure was repeated 100 times for each ligand and the results were clustered using the “gmx cluster” program, part of the GROMACS package (6). The clustering procedure was carried out with a RMSD cut-off of 0.2 nm and the docking poses were not fitted prior to the clustering to capture translational and rotational differences.

VINA affinities and RF-Scores were calculated and the conformational clusters of all ligands were analysed visually in PyMol. Re-docking of compound **1b** (Figure S10A), for which the correct pose was determined experimentally, was used as a control and as a basis for the visual inspection of the remaining compounds. For most molecules displayed in figure S10, visual inspection and the scoring functions were in agreement regarding the most likely pose (exceptions were compound **1c** – Figure S10C, where both scoring functions disagreed with the visually selected conformational cluster, and for compound **2d** – Figure S10F, for which only RF-score and visual inspection were in agreement).

**References**

1.	Di Tommaso P, Moretti S, Xenarios I, Orobitg M, Montanyola A, Chang JM, Taly JF, Notredame C. 2011. T-Coffee: a web server for the multiple sequence alignment of protein and RNA sequences using structural information and homology extension. Nucleic Acids Res 39:W13-7.

2.	Tautz L, Retey J. 2010. A highly convergent synthesis of myristoyl-carba(dethia)-coenzyme A. European J Org Chem 2010:1728-1735.

3.	Morris GM, Huey R, Lindstrom W, Sanner MF, Belew RK, Goodsell DS, Olson AJ. 2009. AutoDock4 and AutoDockTools4: Automated docking with selective receptor flexibility. J Comput Chem 30:2785-91.

4.	Trott O, Olson AJ. 2010. AutoDock Vina: improving the speed and accuracy of docking with a new scoring function, efficient optimization, and multithreading. J Comput Chem 31:455-61.

5.	Wojcikowski M, Zielenkiewicz P, Siedlecki P. 2015. Open Drug Discovery Toolkit (ODDT): a new open-source player in the drug discovery field. J Cheminform 7:26.

6.	Pall S, Abraham MJ, Kutzner C, Hess B, Lindahl E. 2015. Tackling Exascale Software Challenges in Molecular Dynamics Simulations with GROMACS. Solving Software Challenges for Exascale 8759:3-27.
